# Stage-resolved iPSC-to-motoneuron differentiation: Metabolic switch & mitochondrial remodeling

**DOI:** 10.64898/2026.03.25.714145

**Authors:** John Jbeily, Annamarija Raic, Mathias Hafner, Rüdiger Rudolf

## Abstract

Development of motoneurons from stem cells is characterized by a change from glycolytic to oxidative metabolism. Since this transition remains poorly understood, we examined it at five distinct differentiation stages from hiPSC to motoneuron. While a direct comparison of hiPSCs and mature motoneurons confirmed the expected glycolytic-to-oxidative shift, the intermediate stages showed that the conversion was not monotonic. After an initial drop of glycolysis at the hiPSC-to-neuroepithelial transition, late neuroepithelial cells showed intermittent peaks of the glycolytic marker lactate dehydrogenase A and the metabolic regulator TIGAR. Furthermore, the lactate-produced-to-glucose-consumed ratio remained elevated. A fully oxidative phenotype was only assumed upon progress from neural progenitors to motoneurons, portrayed by a definitive drop of the lactate-produced-to-glucose-consumed ratio, an increase of mitochondrial membrane charging, and shifts from lactate dehydrogenase A to B, from pyruvate dehydrogenase to anaplerotic pyruvate carboxylase, and from Mitofusin 1 to 2. Together, our data show that metabolic maturation in human motoneurons does not occur as a simple switch. Instead, it unfolds through distinct stages in a directional yet nonlinear manner.

## Introduction

Lower motoneurons (MNs) are crucial for regulating voluntary muscle contraction. MNs are cholinergic CNS neurons whose cell bodies reside in the ventral horn of the spinal cord while their enormous axons extend over long distances to reach their distal target organs, i.e., skeletal muscles. Both neurodevelopmental as well as adult-onset disorders, including gestational diabetes, embryonic dysgenesis, amyotrophic lateral sclerosis (ALS), and Charcot-Marie-Tooth disease Type 2A, suggest an important role of energy metabolism in their proper functioning (Ornoy *et al*, 2021; Ornoy *et al*, 2015; Vandoorne *et al*, 2018; Mou *et al*, 2021; van Lent *et al*, 2021). During early development, neurons of the CNS differentiate in a multi-step process from pluripotent stem cells over neuroepithelial cells to neuronal precursor cells (NPCs), and ultimately to mature neurons. There is a broad consensus that this developmental path is accompanied by a switch of energy metabolism from glycolytic to oxidative (Danačíková *et al*, 2025; Zheng *et al*, 2016; Agostini *et al*, 2016; Vandoorne *et al*, 2019). This was largely demonstrated using cortical neurons, which were in vitro differentiated from induced pluripotent stem cells (iPSCs). Indeed, proteomic profiling (Danačíková *et al*, 2025) of iPSCs expressing inducible neurogenin 2 (iNGN2) to generate cortical neurons showed already seven days upon neural induction a marked upregulation of neuronal markers as well as enzymes for oxidative metabolism, concomitant with a reduction of glycolytic enzymes. Along these lines, oxygen consumption rate and NAD(P)H measurements yielded functional evidence for an increase in respiratory capacity and metabolic flexibility upon neuronal differentiation. Furthermore, ^13^C-metabolic tracing suggested that neurons may not utilize only glucose as a metabolic substrate to drive mitochondrial respiration (Danačíková *et al*, 2025). These findings were supported by a study using a similar NGN2-based in vitro model, which concentrated on the maturation of neurons from the NPC stage on (Zheng *et al*, 2016). Their transcriptional profiling identified a switch of glycolytic enzyme isoforms typical for aerobic glycolysis (i.e., LDHA and HK2) in NPCs to those more linked to standard metabolism (HK1) or fueling lactate via pyruvate back into the TCA cycle (LDHB). This was corroborated by a reduction of the lactate-produced-to-glucose-consumed ratio (lp/gc) from 1.61 to 0.35, comparing NPCs and neurons (Zheng *et al*, 2016). Notably, there is evidence for mechanistic links between energy metabolism and the progress of neural differentiation. Indeed, overexpression in NPCs of HK2 and LDHA inhibited the progress to the expression of neuronal markers, and under these conditions an increase of external pyruvate as a direct feed into the TCA cycle rescued the halt of neuronal differentiation (Zheng *et al*, 2016). This suggested that a shutoff of glycolysis could be essential for continued differentiation. Similarly, TIGAR, a major negative regulator of glycolysis was found to peak in primary mouse NSCs, and its siRNA-mediated downregulation impaired progression of NSCs to neurons, the timed increase of LDHB expression, and the shift to a mitochondrial, oxidative phenotype (Zhou *et al*, 2019).

Thus, while a couple of studies investigated the metabolic switch in cortical neurons, analogous information on MNs is scarce. Yet, the general feature of a switch from glycolytic to oxidative metabolism in MNs derived from hES and iPSC was previously confirmed by respiration and gene expression analysis (O’Brien *et al*, 2015). Furthermore, iPSCs and MNs of healthy and ALS subjects were investigated by ^13^C-metabolic tracing and oxygen consumption rate measurements (Vandoorne *et al*, 2019). This confirmed that healthy iPSCs were preferentially glycolytic and MNs oxidative. Notably, MNs showed anaplerotic activity, shuttling pyruvate not only via the regular path, i.e., the pyruvate dehydrogenase complex (PDH), but also through pyruvate carboxylase (PC) into the TCA cycle. This might have resulted from an increased lactate usage to generate pyruvate via LDHB. As LDHB also produces NADH (Urbańska & Orzechowski, 2019), which strongly inhibits PDH by mediating the phosphorylation of its A1 subunit (PDHA1) (Patel & Korotchkina, 2006), it sounded reasonable that an alternative route into the TCA cycle was needed; another argument for the need of PC in maturing MNs was to compensate for the loss of the TCA cycle intermediate α-ketoglutarate, being consumed for the production of neurotransmitters in MNs (Vandoorne *et al*, 2019). Although this study did not reveal overt differences between healthy and ALS-derived cells, other reports showed impaired glucose metabolism, mitochondrial respiration, and ATP synthesis in mouse and patient samples from ALS spinal cord (Miyazaki *et al*, 2012; Browne *et al*, 1998; Mattiazzi *et al*, 2002).

Thus, even though there is a general consensus on a switch from glycolysis to oxidative metabolism during early (moto)neuronal development, the studies so far have only picked two or three developmental stages (typically iPSC or NPC and neuron) and they have mostly used glucose- or lactate-centered metabolite tracing experiments, oxygen consumption rate analysis, and transcriptomics/proteomics as readouts. We were interested to learn if the glycolytic-to-oxidative switch is a linear process, if different components of the metabolism (e.g., glycolysis exit, TCA cycle entry, mitochondrial maturation and activity) would switch simultaneously or in sequence, and how metabolites would shuttle between cells and their environment during early neuronal development. Therefore, we investigated the interplay between glycolytic and oxidative metabolism at five different stages of in vitro MN development, i.e., iPSC, early and late neuroepithelial progenitors, NPC, and MN. Immunofluorescence-based analysis focused on TIGAR as a potential key glycolytic regulator, on the LDH complex as the exit from glycolysis, on PDH and PC as entry routes into the TCA cycle, and Mfn1/2 and Drp1 as major components of the mitochondrial fusion/fission machinery. Mitochondrial morphology and membrane potential were examined by live cell imaging and subsequent computational analysis to retrieve a functional and morphological correlate of the metabolic processes in action. Finally, NMR-based extracellular metabolite analysis was used to relate the immunofluorescence and mitochondrial data to the metabolic activity of the cells in an untargeted manner. This revealed that, (i) while iPSC and MN stages were characterized by clear glycolytic and oxidative profiles, respectively, the intermediate stages showed mixed phenotypes that were not unidirectionally and linearly moving towards oxidative; in particular TIGAR, LDHA, and LDHB showed remarkable variation during the developmental profile; (ii) different components of the metabolic machinery switched at distinct time points, with the mitochondrial potential rising already at the neuroepithelial stage but a complete switch only occurring at the MN stage; (iii) while pyruvate was steadily consumed throughout differentiation, iPSCs showed a peculiar signature for amino acids and two further small metabolites, i.e., release of glutamine, glycine, choline, and acetone into the extracellular medium; (iv) upon transition from precursor to maturing MNs, phosphorylation-dependent inhibition of PDH coincided with a marked increase in LDHB, Mfn2/Mfn1 ratio, mitochondrial membrane potential (ΔΨ_m_), and PC. These findings suggest that iPSC-derived MNs strongly rely on alternative oxidative metabolic routes, such as the use of external lactate and pyruvate in combination with anaplerotic TCA cycle entry. Our work provides a longitudinal and multimodal analysis of early human motoneuronal development suggesting a stepwise and modular metabolic rewiring that links switches of glycolysis exit, TCA cycle entry, mitochondrial remodeling, and crosstalk with the extracellular milieu.

## Results

### Expression profiles of metabolic key enzymes suggest a non-linear conversion of from glycolytic to oxidative and increasing anaplerosis during motoneuron development

By directly comparing hiPSC and MNs using metabolic tracing and oxygen consumption rate analysis, a previous study had found a switch of metabolism from glycolytic to oxidative and the presence of anaplerosis upon differentiation to MNs (Vandoorne *et al*, 2019). To complement these data with other techniques and to learn if the process of metabolic conversion was steady and unidirectional, we performed a set of investigations at five different developmental stages from hiPSC to MN. In detail, the following stages of a previously characterized small-molecule-based differentiation protocol (Hörner *et al*, 2021; Du *et al*, 2015) were included: hiPSC (d0), early (d3) and late neuroepithelial progenitor (d8), NPC (d14), and maturating MN (d25). The proper differentiation trajectory was approved by a small marker panel ranging from Oct-4 and Sox-2 for d0, Olig-2 for d14, and Isl-1, Hb-9, and Tuj1 for d25 (Fig. S1). The first part of investigating a putative metabolic switch along these five differentiation stages employed an expression analysis of key metabolic enzymes using immunofluorescence. This included antibodies against the LDH complex to probe the exit of glycolysis, TIGAR as a negative regulator of glycolysis and gatekeeper of neuronal differentiation, PDHA1 and its inactive phosphorylated version, as well as PC as different entries into the TCA cycle (see Fig. S2 for an overview of markers and metabolites addressed in the study). Although this panel corroborated a clear switch from a glycolytic to an oxidative profile upon direct comparison of hiPSCs (d0) and MNs (d25), incorporation of the data taken at intermediate stages (d3, d8, d14) did not suggest a linear and unidirectional conversion route (Fig. 1). First, pan-LDH and LDHA decreased overall from d0 to d25, but both showed an intermittent peak at d8. Second, although LDHB increased from d0 to d25, it exhibited significant drops at d3 and d14 (Fig. 1). Third, relative TIGAR expression was steady for most time points but, again, d8 stood out with a peak. Fourth, pan-PDHA1 displayed a rising trend on d8 and then dropped on d14 and d25. This was mirrored by a relative drop of the inactive phosphorylated PDHA1 (pPDHA1) from d0 to d8 and a rising trend from d14 to d25. Lastly, PC showed a similar expression profile as LDHB with an overall increase from d0 to d25 and intermediate drops on d3 and d14. In summary, while the expression profiles of key metabolic markers corroborated earlier work stating a neurodevelopmental conversion of energy metabolism from glycolytic at the stem cell stage to oxidative at the neuron stage, they also indicated a non-linear trajectory of the rewiring. In particular, d8 cells showed a peculiar mixed phenotype between glycolytic and oxidative. Furthermore, the anaplerotic signature of an increased expression of PC at the expense of active PHDA1 became visible at d14 and even more so at d25.

**Figure 1:**
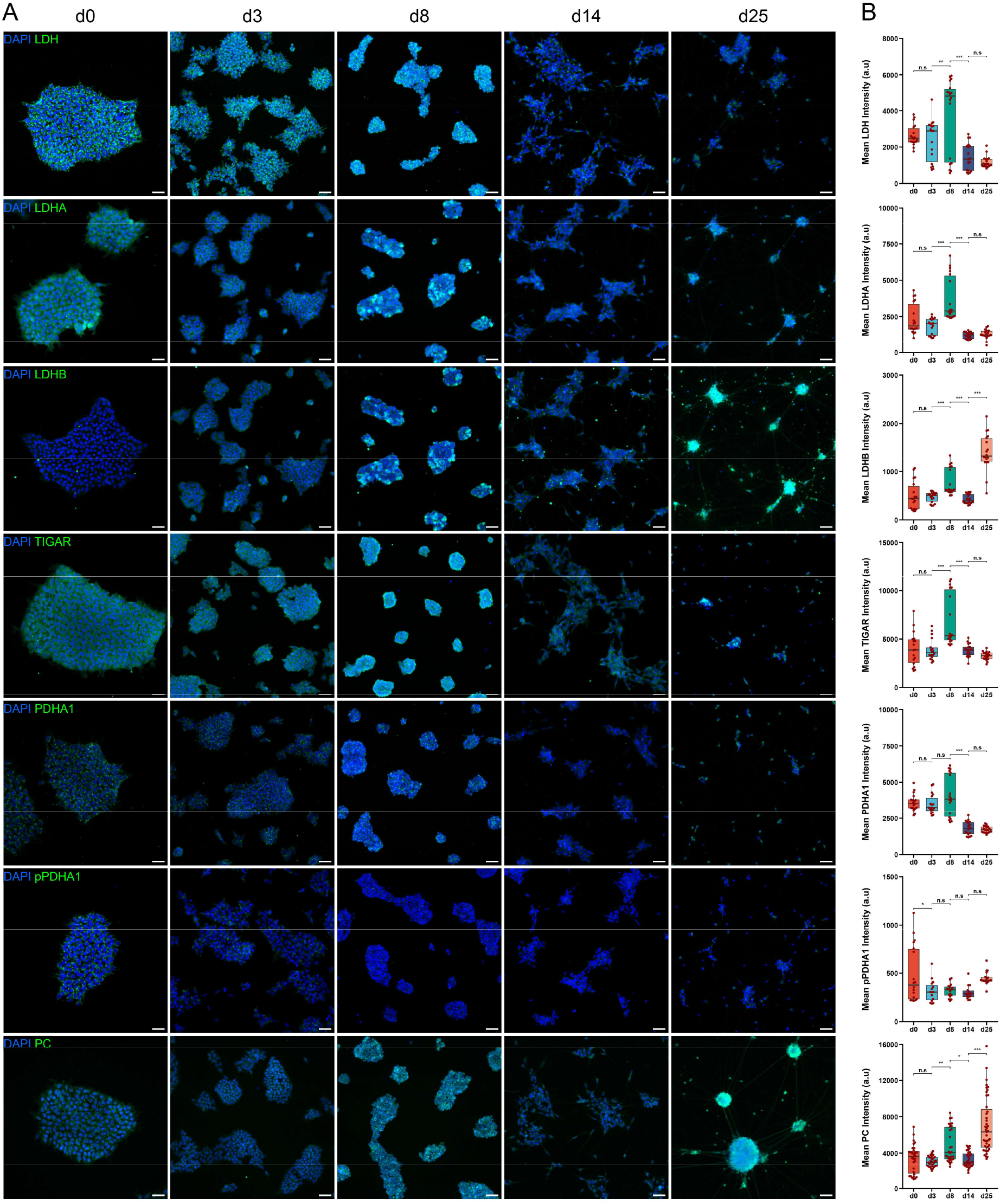
A non-monotonous transition from glycolysis to oxidative metabolism and increasing relevance of anaplerosis are suggested by key marker expression profiles. hiPSCs were differentiated into MNs and fixed on days 0, 3, 8, 14, and 25 of the differentiation protocol. Then, IF was executed using DAPI to stain nuclei and antibodies against markers as indicated in the figure panels. Widefield microscopy and image analysis were performed. (A) Representative widefield micrographs taken at developmental days as indicated on top (d0-d25). Images depict DAPI staining in blue and antibody staining in green. Scale bars, 50 µm. (B) Box/Whisker plots show quantitative IF analysis of antibody staining mean intensities per visual field corresponding to the data shown in A. Significance was measured using either parametric or non-parametric tests, depending on the normality of data distribution. n.s, not significant; * p < 0.05; ** p < 0.01; *** p < 0.001.

### The transitions from stem cell to neuroepithelial and from neuroepithelial to neural progenitor feature a stepwise increase of mitochondrial membrane potential

The expression profiles of key metabolic enzymes suggested a non-linear, stepwise conversion from glycolytic to oxidative. Next, we addressed if this was reflected by ΔΨ_m_ as a proxy of their metabolic activity (Tovar-Ferrero *et al*, 2025). To do so, live imaging of cells at the five different developmental stages (d0, d3, d8, d14, d25) was performed using the ΔΨ_m_ indicator dye, JC1. Due to its cationic nature, JC1 accumulates in mitochondria with high membrane potential. In doing so, it shifts its fluorescence spectrum from green to red (Glaß *et al*, 2020; Overly *et al*, 1996). Thus, cells were labeled with JC1 and then visualized with confocal live cell imaging, simultaneously recording green and red fluorescence channels. As depicted in Fig. 2A, all cells nicely took up the dye and most of it was located in mitochondrial structures. Notably, while red JC1 signal was hardly visible at d0 (Fig. 2A), it became stronger at the following time points and the red/green JC1 ratio increased more and more from d3 to d25 (Fig. 2B). Quantitative JC1 assays normally involve determining both the area and intensity ratios of red/green JC1 signals. While the first is a proxy for the proportion of charged mitochondria of the total mitochondrial network, the second indicates the overall mitochondrial polarization state. Both area ratio and intensity ratio significantly increased from d0 to d3 (Fig. 2C and D). While after d3 the area ratio showed only a trend to further increase, particularly between d8 and d14 (Fig. 2C), a second significant rise between d8 and d14 was evident for the intensity ratio (Fig. 2D). These data suggested, that large parts of the mitochondrial network started to polarize already at d3, but that it reached its maximal polarization only as of d14. Since at d25 a wide network of neurites was present, it was also possible to segregate intensity ratio values from cell bodies vs. neurites. This revealed significantly higher ratios in neurites (Fig. 2E), suggesting a differential metabolic activity in both cellular compartments. In summary, live cell imaging of JC1 red/green ratio values suggested an increase of mitochondrial activity in two steps, i.e., from d0 to d3 and from d8 to d14.

**Figure 2:**
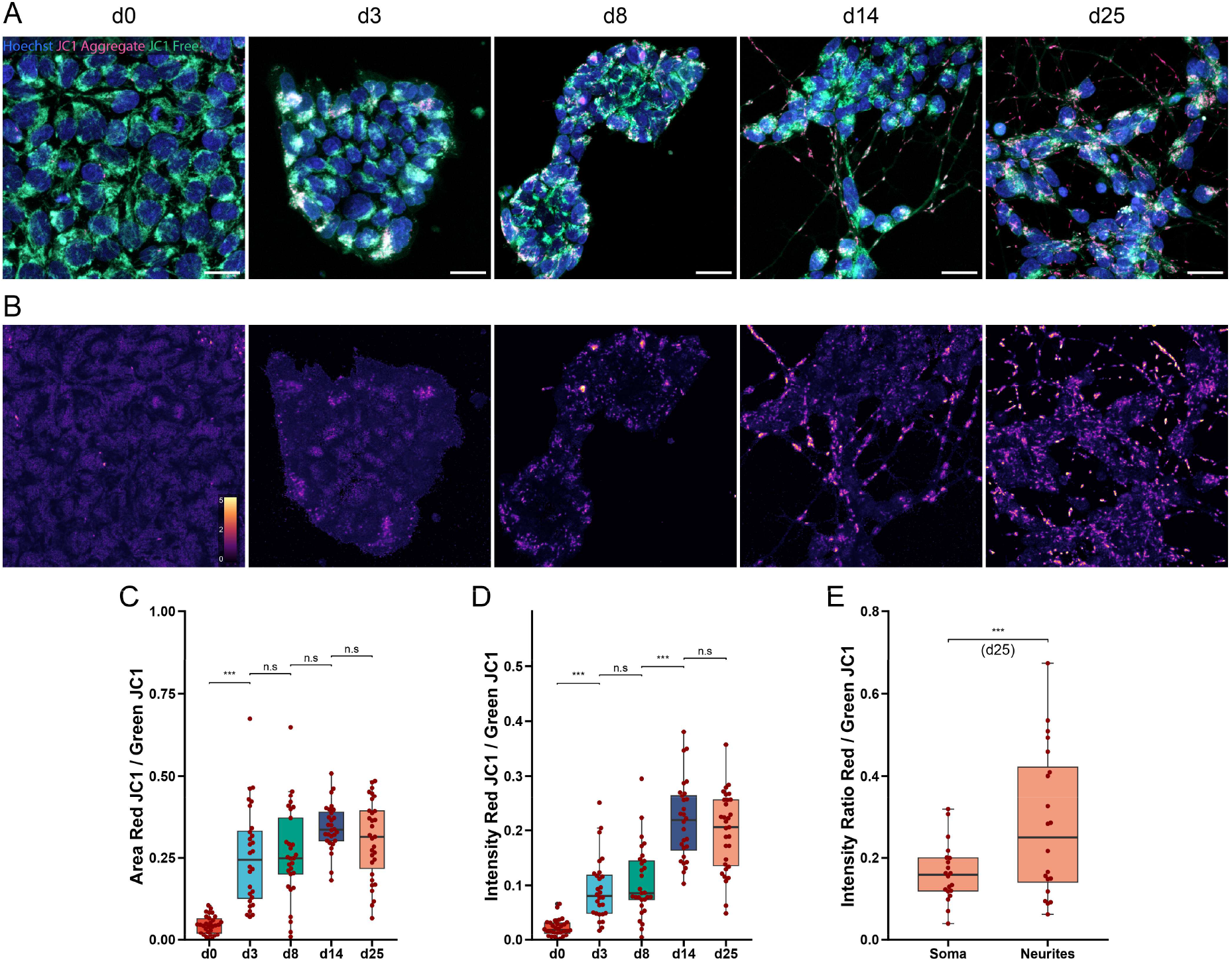
Mitochondrial membrane potential rises in two steps from the iPSC-to-neuroepithelial and from the neuroepithelial-to-NPC transition. hiPSCs were differentiated into MNs and live imaged as whole-cell z-stacks with Hoechst and JC-1 dyes for nuclei and mitochondrial membrane potential, respectively, on days 0, 3, 8, 14, and 25 of the differentiation protocol. Image analysis was performed. (A) Representative confocal micrographs taken at developmental days as indicated on top (d0-d25). Images depict Hoechst staining in blue, aggregated JC1 in red, free JC1 in green. Scale bars, 20 µm. (B) Pseudo-colored images show JC1 aggregated/free ratios with high and low values in purple and orange/white, respectively. Images correspond to those in A. Pseudo-color scale bar in left image. (C-E) Box/Whisker plots show quantitative analysis of data sets as exemplified in A as a function of differentiation time for area ratio JC1 aggregated/free (C) and intensity ratio JC1 aggregated/free in the whole image (D) as well as segregated values for cell bodies and neurites (only d25, E). Significance was measured using the Kruskal-Wallis non-parametric test with a Wilcoxon post hoc test. n.s, not significant; * p < 0.05; ** p < 0.01; *** p < 0.001.

### Motoneuronal differentiation is accompanied by increasing mitochondrial compaction, number, and Mfn2/Mfn1 and Drp1 levels

Next to membrane potential, also mitochondrial morphology is considered to reflect their metabolic activity (Galloway *et al*, 2012; Scandella *et al*, 2023). Thus, we aimed to characterize the mitochondrial networks and the corresponding fusion/fission machineries from d0 to d25. To gain insight into the morphology of mitochondria, cells were stained with Mitotracker Red CMXRos and were then live-imaged in 3D using confocal microscopy with voxel sizes of 80 x 80 x 300 nm. Notably, we opted for live cell imaging, because mitochondrial morphology was strongly impaired by PFA fixation (Fig. S3 and Text S1). An Ilastik-based 3D-segmentation model was trained and applied to segment the live cell Mitotracker signals in 3D. As illustrated in Fig. 3A, all cells showed a nicely formed mitochondrial network, that extended throughout the cell bodies at d0 and appeared to be increasingly clustered next to the nuclei with progressing differentiation. The automated 3D-segmentation was robust (see Fig. 3B for a less dense region of d25 motoneuronal neurites to exemplify the segmentation accuracy, and the supplementary video for a 3D rendering of the segmentation masks) and yielded large but discrete mitochondrial entities (Fig. 3A, lower row, different colors), which appeared more and more compacted from d0 to d25 (Fig. 3A, lower row). A quantitative analysis revealed a decreasing radial distance of mitochondria from the mitochondrial network center of mass (Fig. 3C), corroborating the visual impression of mitochondrial compaction. This was particularly prominent from d0 to d8 and then remained relatively stable. The number of mitochondria per cell (Fig. 3D) and the average total volume of mitochondria per cell (Fig. 3E) showed a largely complementary behavior: while d0, d3, and d14 cells displayed the lowest number and the highest volume of mitochondria, d25 neurons had the largest number of mitochondria per cell but with the lowest total volume. To approach mitochondrial branching/complexity, the fractal dimension of the mitochondrial 3D-segments was determined. This describes how completely a given pattern fills space. Higher fractal dimensions mean more complexity and finer details. This showed the highest values on d0 and d8, and significant drops between d0 and d3 as well as between d14 and d25 (Fig. 3F). In summary, during motoneuronal differentiation from hiPSC, the mitochondrial network showed increased compaction and mitochondria number, a decrease in mitochondrial volume, and an intermediate complexity. Again, these trends were not developing in a monotonous but rather contorted manner, with d8 reflecting intermediate characteristics between hiPSC and neuronal features. Next, the expression profiles of major mitochondrial fusion/fission machinery components were determined to accompany the morphological data. In detail, Mfn1 and Mfn2 of the fusion apparatus as well as the fission marker Drp1 were stained. (Fig 3G) All of these showed clearcut but distinct profiles. While Mfn1 was strongly expressed at early time points, peaked at d8, and was almost completely absent at d25, Mfn2 was low at d0 and d3, showed a first intermediate peak at d8, and its maximum at d25. (Fig 3H). Finally, Drp1 increased from d0 to d3, decreased on d14, and then peaked, like Mfn2, on d25. Thus, whereas all three components were relatively high at d8, suggesting an ample remodeling of the mitochondrial network at this stage, differentiated neurons displayed a high level of Drp1 and a strong switch from Mfn1 to Mfn2 (Fig 3G, H).

**Figure 3:**
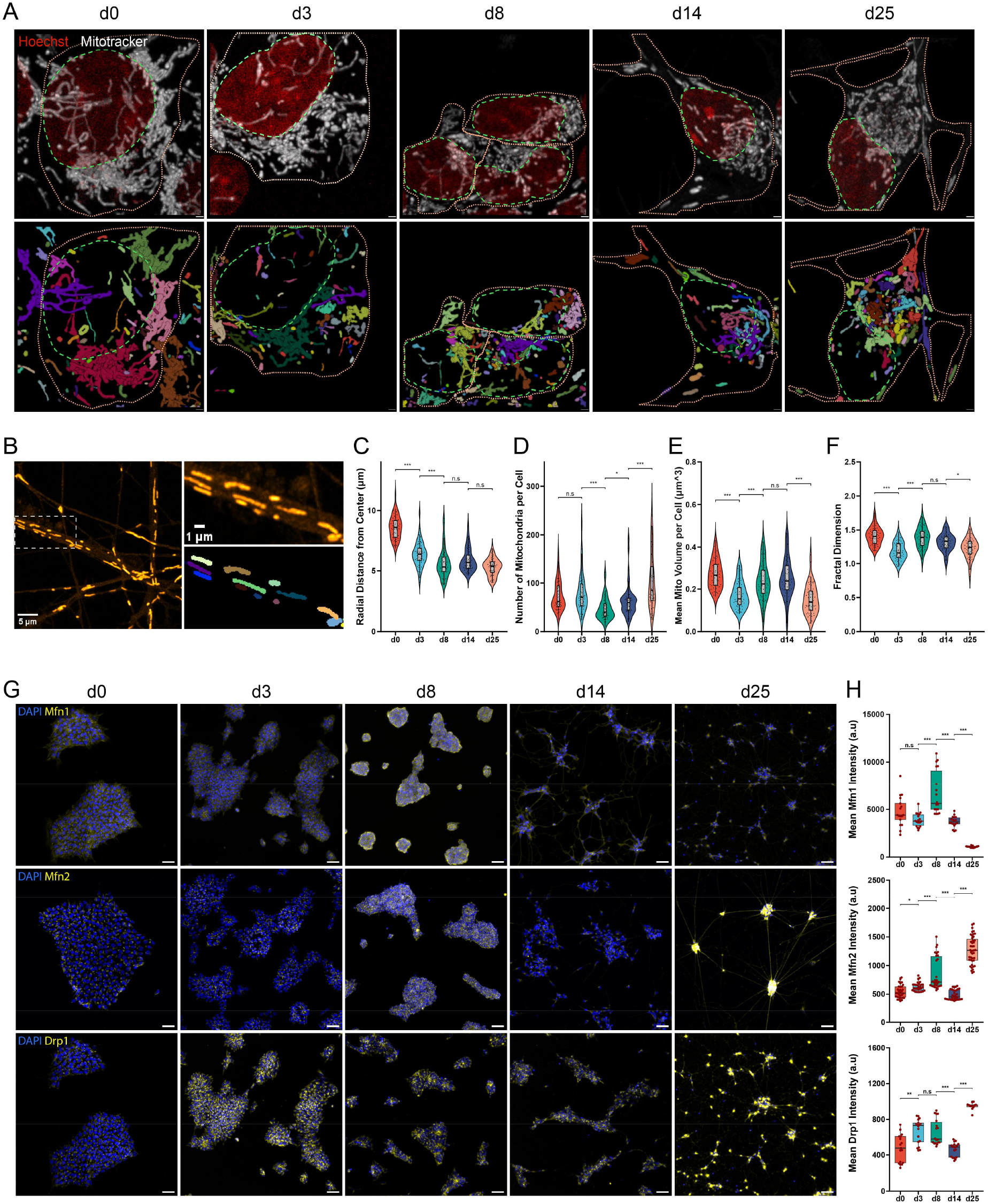
Mitochondria compact during hiPSC-to-motor neuron differentiation, with stage-dependent expression of the key fission/fusion markers Mfn1, Mfn2, and Drp1. hiPSCs were differentiated into MNs and live imaged as whole-cell z-stacks with Hoechst and Mitotracker dyes for nuclei and mitochondrial morphology, respectively, on days 0, 3, 8, 14, and 25 of the differentiation protocol. Image analysis was performed. (A) Representative confocal micrographs taken at developmental days as indicated on top (d0-d25). Images depict Hoechst staining in red and Mitotracker in gray. Scale bars, 1 µm. Green dashes outline the nucleus border, while orange dashes outline the cell border. Lower panels show segmented mitochondrial objects within the same cells. Colored labels denote individual mitochondrial networks. (B) A representative confocal micrograph of small single mitochondria within motor neuronal (d25) neurites with the right panel showing a zoom and relevant segmentation masks. (C-F) Violin plots show quantitative analysis of data sets as exemplified in A as a function of differentiation time for radial distance from the center of mass of all the mitochondria clusters per cell (C), number of mitochondria per cell (D), mean mitochondrial volume per cell (E), and fractal dimension (F), a proxy for reporting the extent of mitochondrial branching. Each data point corresponds to a pooled data set of all the single mitochondrial networks inside a single cell. 50 distinct cells were analyzed for every time point. Significance was measured using the Kruskal-Wallis non-parametric test with a Wilcoxon post hoc test. n.s, not significant; * p < 0.05; ** p < 0.01; *** p < 0.001. (G) Representative widefield micrographs taken at developmental days as indicated on top (d0-d25). Images depict DAPI staining in blue and antibody staining in yellow. Scale bars, 50 µm. (H) Box/Whisker plots show quantitative IF analysis of antibody staining mean intensities per visual field corresponding to the data shown in G. Significance was measured using either parametric or non-parametric tests, depending on the normality of data distribution. * n.s, not significant; * p < 0.05; ** p < 0.01; *** p < 0.001.

### Metabolite profiling confirms the stem cell and neuron stages to be particularly different during motoneuron development

To set the enzymatic and mitochondrial profiles into a metabolite context, an untargeted NMR-based metabolomic analysis of cell supernatants was performed for all five differentiation time points from d0 to d25. This yielded clear peaks for eight amino acids, three small metabolites, and lactate (Fig. 4A). Furthermore, glucose concentrations were determined using a photometric assay. For the measurements, cells at the five indicated differentiation points were incubated for 24 h in fresh medium, and the media were then analyzed and referenced to the same media incubated for 24 h in the absence of cells. Thus, release into or consumption of the metabolites from the media were assessed. Regarding the amino acids (Fig. 4B), quantitative analysis showed that the non-essential glutamine and glycine, which are involved in nitrogen transfer, glutathione production, and DNA synthesis (DeBerardinis & Cheng, 2010; Cruzat *et al*, 2018; Locasale, 2013), were released at d0. Indeed, glutamine and glycine were present in the extracellular supernatant at concentrations of 3.08 mM ± 0.11 mM and 0.44 mM ± 0.16 mM, respectively, above reference at d0. At the other time points, they were oscillating around zero, except for glutamine at d8, being at 0.35 mM ± 0.16 mM. Conversely, the essential branched-chain amino acids valine, leucine, and isoleucine were consumed at d0 and were close to zero at all other time points. Finally, the aromatic amino acids tyrosine and phenylalanine as well as the essential histidine were always close to zero. Concerning the small metabolites choline and acetone (Fig. 4C), these were released at d0 and consumed at all other time points, while the lactate/TCA donor pyruvate was consumed at all time points, with a maximum at d0 (-0.20 mM ± 0.04 mM, Fig. 4C). Thus, for amino acids and three small metabolites, the only time point with a peculiar pattern was d0. This was different for lactate production and glucose consumption (Fig. 4D). Although both were highest at d0 and then continuously decreased for the entire remaining observation period, the lp/gc, commonly used as a measure of the efficiency of oxidative metabolism (Teslaa & Teitell, 2014; Orlovskiy *et al*, 2025), showed two significant drops, i.e., between d0 and d3 as well as between d14 and d25. This suggested the presence of at least two major steps of metabolic rewiring from glycolytic to oxidative.

**Figure 4:**
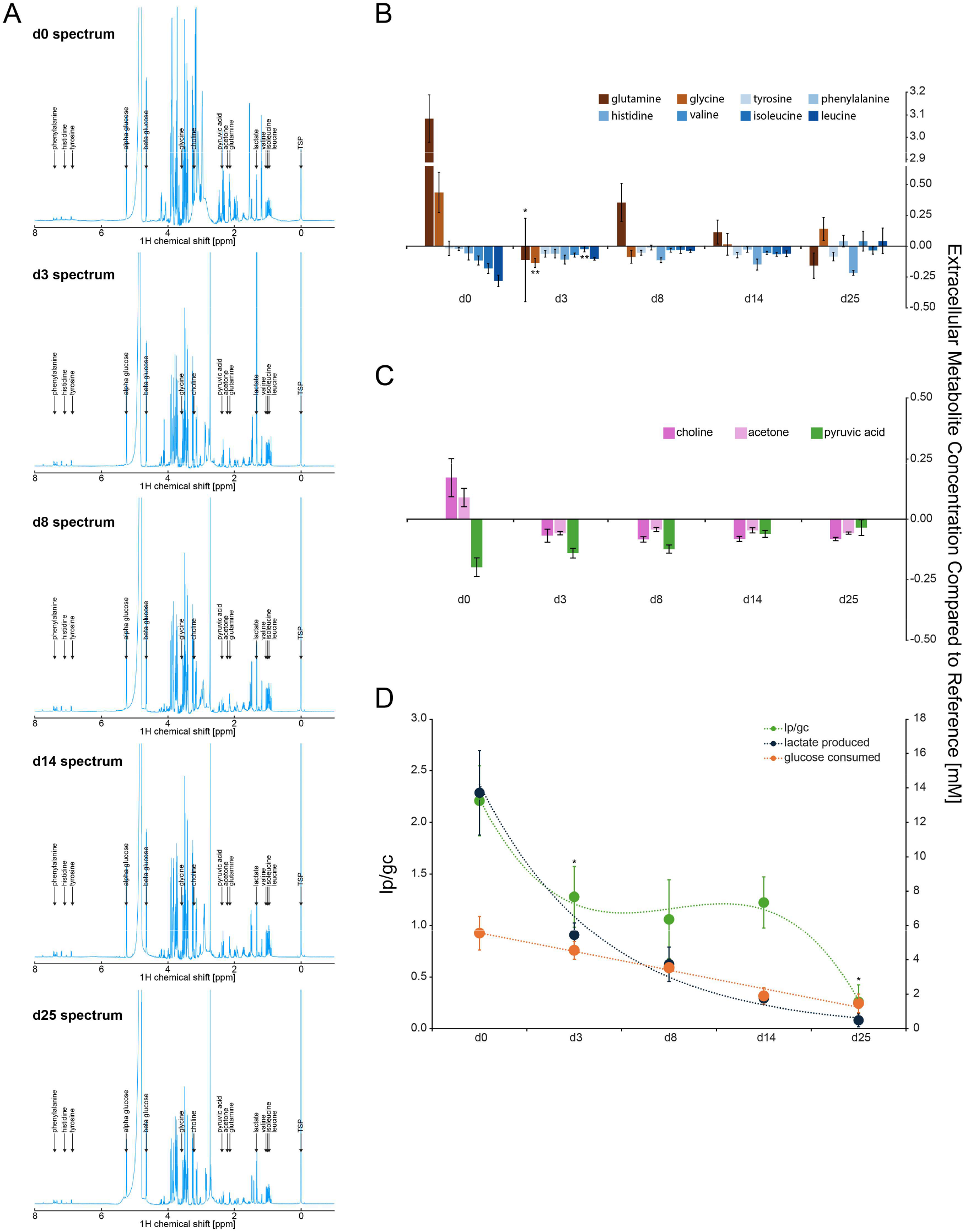
NMR-based analysis of supernatants suggests an inefficient metabolism of amino acids, choline, and keto bodies at d0, and corroborates a non-linear transition to oxidative metabolism. hiPSCs were differentiated into MNs and supernatants were taken on days 0, 3, 8, 14, and 25 of the differentiation protocol. Then, supernatants and corresponding reference medium samples were analyzed by NMR, followed by spectral analysis. Glucose values were determined separately. (A) Representative NMR-spectra of media and supernatants taken at developmental days as indicated on top (d0-d25). (B-C) Bar graphs depict concentrations of metabolites as indicated in the supernatant taken at time points as indicated (d0-d25). Mean ± SEM; all values were referenced to corresponding media. Significance was measured using either parametric or non-parametric tests, depending on the normality of data distribution. * n.s, not significant; * p < 0.05; ** p < 0.01; *** p < 0.001. (D) Line graph depicts lactate produced (dark blue), glucose consumed (orange), and their ratio (green) at time points as indicated (d0-d25). Mean ± SEM; all values were referenced to corresponding media. Consecutive time points were compared using one-tailed Welch’s t-tests. * n.s, not significant; * p < 0.05; ** p < 0.01; *** p < 0.001.

## Discussion

Neurons are among the most energy-demanding cell types in the human body, requiring finely tuned bioenergetic programs to sustain development, synaptic activity, and long-term survival (Iwata *et al*, 2023; Magistretti & Allaman, 2015). Their energy requirements are extraordinary: although the brain constitutes only about 2 % of body mass, it consumes nearly 20 % of resting metabolic energy, primarily to support neuronal signaling (Attwell & Laughlin, 2001). Glucose metabolism is central to meeting these needs, serving both as the primary source of adenosine trisphosphate (ATP) and as a provider of biosynthetic intermediates required for proliferation, differentiation, and neurotransmitter synthesis (DeBerardinis & Thompson, 2012; Vander Heiden *et al*, 2009). During neuronal differentiation, i.e., the transformation of a pluripotent stem cell into a mature neuron, the cell undergoes a profound metabolic reorganization, transitioning from a glycolytic profile toward increased reliance on oxidative phosphorylation (Agostini *et al*, 2016; Beckervordersandforth *et al*, 2017; Rumpf *et al*, 2023). This shift normally reflects both expansion of mitochondrial networks and changes in mitochondrial activity. While the developmental shift from glycolysis to oxidative phosphorylation is well established in neurons of the brain, the evidence in MNs is still scarce. Furthermore, the precise mechanism controlling the entry of pyruvate into the TCA cycle remains poorly understood. Canonically, it has been shown that the PDH complex is the central gatekeeper converting pyruvate into acetyl-CoA and thereby linking glycolysis to mitochondrial oxidation (Stacpoole & McCall, 2023; Wieland, 1983; Kim *et al*, 2006; Kaplon *et al*, 2013) (see Fig. S2 for overview). PDH activity is strongly regulated by reversible phosphorylation, which inhibits oxidative decarboxylation of pyruvate and reduces acetyl-CoA availability (Patel & Korotchkina, 2006; Gray *et al*, 2014; Korotchkina & Patel, 2001; Bowker-Kinley *et al*, 1998; Huang *et al*, 1998). In many cell types, PDH activation is concomitant with increased oxidative metabolism. In MNs, however, this regulation had not been explored. One study showed that induced pluripotent stem cell (iPSC)-derived motor neurons upregulated pyruvate carboxylase (PC) as a pathway to shuttle carbon into the TCA cycle independently of PDH (Vandoorne *et al*, 2019). In addition, PC was found to play a role in maintaining neuronal anaplerosis and supporting neurotransmitter synthesis (Mason *et al*, 2007; Rothman *et al*, 2011). Taken together, this grants plausibility to the idea that developing MNs may, in fact, recruit non-canonical pathways to shuttle carbon into the TCA cycle, ensuring sufficient anaplerotic support for growth and maturation. Whether or not PDH activity is altered, has not yet been clearly addressed.

Our data support the glycolytic-to-oxidative switch paradigm during iPSC-to-MN differentiation (Vandoorne *et al*, 2019), but they argue against a simple linear progression. Instead, the transition occurred in distinct stages with partial flips and reversals of discrete metabolic modules, such as the glycolysis exit, the mitochondrial charging, and the TCA cycle entry. Indeed, while some parameters, including lactate production, glucose consumption, pyruvate consumption (Fig. 4C, D), and nuclear size (Fig. S4), changed monotonically from d0 to d25, consistent with a gradual egress of glycolysis and proliferative stem-cell-like metabolism, total LDH and LDHA flipped from average at d0 to high at d8 to low at d25 (Fig. 5A). Similarly, the lp/gc ratio and ΔΨ_m_, represented by JC1 Int R/G, although uniform in direction, with lp/gc decreasing and ΔΨ_m_ increasing, underwent changes at discrete phases of differentiation (Fig. 5A). In contrast, several metabolic enzymes and mitochondrial-dynamics markers showed irregular or biphasic behavior. This is important, because a simple comparison between hiPSCs and mature MNs would suggest a binary switch, whereas the intermediate stages reveal a more structured process. In that sense, our data extend previous work on neuronal metabolic reprogramming in the context of neuronal (Agostini *et al*, 2016; Zheng *et al*, 2016) and motoneuronal (Vandoorne *et al*, 2019; O’Brien *et al*, 2015) differentiation by showing how this conversion develops over time.

**Figure 5:**
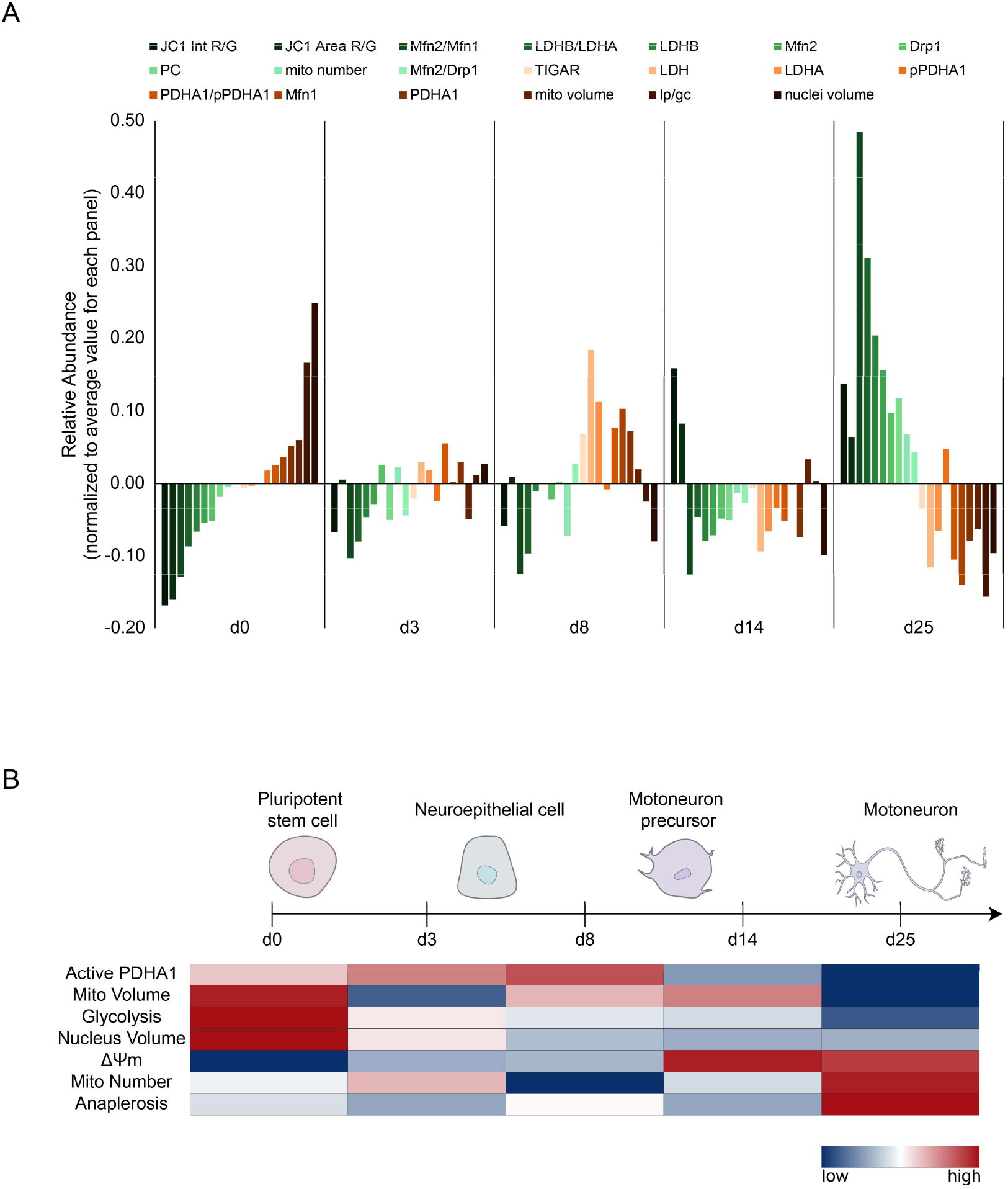
Integrated metabolic and mitochondrial marker profiling reveals stage-dependent, non-monotonic remodeling across iPSC-to–motor neuron differentiation. Data collected across differentiation time points (d0, d3, d8, d14, d25) were compiled from the assays shown throughout this study and summarized as a multi-marker overview. (A) Bar plot showing the relative abundance of key metabolic, mitochondrial, and morphological features at each differentiation stage (including JC-1 red/green ratio metrics, LDH/LDHA/LDHB, TIGAR, PDHA1 and pPDHA1, and mitochondrial/nuclear morphology readouts). Functionally informative ratios were additionally computed for meaningful comparisons, such as PDHA1/pPDHA1 for canonical TCA entry inhibition, Mfn2/Mfn1 for fusion regulator balance, LDHB/LDHA for glycolysis exit, and Mfn2/Drp1 for fusion-fission balance. Each feature was normalized within-feature across time points by mean-centering and sum-scaling; values, therefore, reflect relative deviations across differentiation. (B) Scheme representing the most prominent changes observed throughout differentiation at timepoints as indicated. In the heatmap, Dark blue indicates low relative abundance of a feature, and dark red indicates its high relative abundance. Colors were determined by directly comparing the values shown in A within the same feature.

The first major shift occurred between d0 and d3, i.e., from iPSC to early neuroepithelial stage. This was marked by a first drop in lp/gc, the onset of mitochondrial polarization, and the loss of the d0-specific release of glutamine, glycine, choline, and acetone into the extracellular space. Together, these changes suggest that the cells had already started to leave the pluripotent metabolic program very early during differentiation. That fits well with the broader view that pluripotent cells rely strongly on glycolysis not only for ATP production, but also to sustain biosynthetic fluxes associated with growth and self-renewal (DeBerardinis & Thompson, 2012; Vander Heiden *et al*, 2009). It also agrees with previous studies showing that neuronal differentiation is accompanied by early mitochondrial engagement and reduced glycolytic dependence (Danačíková *et al*, 2025; Zheng *et al*, 2016; Agostini *et al*, 2016; O’Brien *et al*, 2015; Beckervordersandforth *et al*, 2017).

Interestingly, d8 was not simply a midpoint on a straight trajectory between hiPSCs and mature MNs, but appeared as a distinct transitional state. LDH and LDHA actually increased compared to hiPSC, consistent with a strong reliance on glycolytic and lactate-linked pyruvate metabolism, a feature typically associated with biosynthetically active cell states (DeBerardinis & Thompson, 2012; Vander Heiden *et al*, 2009). At the same time, mitochondrial polarization had already begun, whereas mitochondrial number remained low and lp/gc did not decrease further, suggesting that mitochondrial activation preceded full oxidative rewiring. The peak in TIGAR at d8 is also notable in this context, since this metabolic regulator enzyme has been found to be essential to neural stem cell differentiation and neuronal fate control through acetyl-CoA-dependent chromatin regulation (Zhou *et al*, 2019). The concurrent elevation of Mfn1, Mfn2, and Drp1 further argues that this stage was characterized by active mitochondrial remodeling, in line with the known role of fusion and fission in neuronal development and mitochondrial redistribution (Chen *et al*, 2007; Sheng & Cai, 2012; Lee *et al*, 2012; Casellas-Díaz *et al*, 2021; Vevea & Chapman, 2023). Together, these readouts place cells at d8 in a metabolically mixed state, where glycolytic features were still present even though mitochondrial adaptation and differentiation-associated metabolic programming were already underway.

Moving on from d8 to d14, LDH and LDHA fell sharply, suggesting that the previously still active glycolytic program was being downregulated, parallel to what other studies in neuronal differentiation have observed (Zheng *et al*, 2016; Agostini *et al*, 2016). At the same time, ΔΨ_m_ increased strongly, especially at the level of overall polarization, implying that the mitochondrial network was becoming functionally engaged on a broader scale. This corroborates the idea that neuronal differentiation requires early mitochondrial activation before full terminal maturation is reached (Zheng *et al*, 2016; O’Brien *et al*, 2015). Although mitochondria were now more polarized, the lp/gc ratio had not yet decreased further. This hints that mitochondrial charging and oxidative flux are not necessarily equivalent. Yes, a higher ΔΨ_m_ means that mitochondria are increasingly capable of sustaining proton motive force and ATP-linked respiration, but as previously reported, does not always mean an increased mitochondrial activity (Brand & Nicholls, 2011). In our system, this interpretation is also supported by the fact that the PDHA1/pPDHA1 ratio was low at d14, arguing for an inefficient canonical entry of pyruvate into the TCA cycle. Furthermore, also PC and LDHB were still low at this stage, suggesting a still low rate of anaplerosis that could have filled the gap of fuelling the TCA cycle via the lactate-pyruvate-oxaloacetate bridge (see Fig. S2 for an overview). In other words, the machinery for oxidative metabolism was being engaged, but the broader metabolic architecture of the cell still appeared incomplete. This distinction is mechanistically plausible. In differentiating neurons, oxidative maturation is not just a matter of turning mitochondria “on”. It also depends on how many mitochondria are present, where they are positioned, and how their network is remodeled to match the emerging cellular architecture (Sheng & Cai, 2012; Casellas-Díaz *et al*, 2021; Vevea & Chapman, 2023). Our morphology data confirm this view. By d14, the mitochondrial network had already become more compact than at d0, but the increase in mitochondrial number that we observed at d25 had not yet occurred. The high levels of Mfn1, Mfn2, and Drp1 at d8 suggested that this timepoint was dominated by active network remodeling, which may be a prerequisite for the later establishment of a fully neuron-adapted oxidative phenotype. Taken together, the d8-to-d14 interval likely marked the point at which mitochondria became functionally competent, while full metabolic commitment still lagged behind.

The late transition from d14 to d25 looked like the point at which metabolic rewiring was consolidated into a definitive oxidative profile. This phase was marked by the second drop in lp/gc, maximal mitochondrial polarization, an increase in mitochondrial number, and the strongest rise in LDHB, PC, PDHA1/pPDHA1 ratio, Mfn2/Mfn1 ratio, and Drp1. Taken together, these changes argue that late MN maturation is not simply a matter of “more OXPHOS” but of building a mitochondrial system that is decidedly oxidative and neuron-adapted. The higher JC1 ratio in neurites than in soma at d25 supports this, because mature neurons depend on highly functional mitochondria in distal compartments where ATP demand, Ca^2+^ buffering, and local organelle turnover are especially high (Sheng & Cai, 2012; Vevea & Chapman, 2023). Our morphology data point in the same direction: across differentiation, mitochondria became more compact, more numerous, and individually smaller, albeit non-monotonically. In a MN, that kind of remodeling is unlikely to be incidental. Mitochondrial shape, size, and distribution are tightly linked to trafficking, local energy supply, and quality control in polarized neuronal processes (Sheng & Cai, 2012). Mfn2 is particularly relevant here because it contributes not only to mitochondrial outer-membrane fusion, but also to axonal mitochondrial transport, neurite development, and ER-mitochondria coupling, the latter being important for Ca^2+^ transfer and bioenergetic regulation (Mou *et al*, 2021; Casellas-Díaz *et al*, 2021; Brito & Scorrano, 2008; Filadi *et al*, 2015). The late rise in Mfn2, therefore, fits well with the emergence of a more mature neuronal mitochondrial network. The increase in Drp1 should not be read simply as fragmentation or stress, either. In neurons, fission is part of normal network maintenance: it helps generating mitochondria that can be delivered into neurites and axons and supports local turnover of damaged organelles (Kiryu-Seo *et al*, 2016; Zerihun *et al*, 2023). Seen together, the shift from early Mfn1 predominance to late Mfn2 and Drp1 enrichment suggests that mitochondrial remodeling and metabolic maturation go hand in hand. Traditionally, PDH is thought to be the canonical gatekeeper linking glycolysis to mitochondrial oxidation (Patel & Korotchkina, 2006; Stacpoole & McCall, 2023). However, our late-stage profile does not fit a simple model of progressive PDH activation as oxidative potential increases. Pan-PDHA1 rose at d8, then declined again at d14 and d25, while its phosphorylated, and thus inhibited counterpart, pPDHA1, showed the opposite late pattern, increasing from d14 to d25, even if statistically insignificant. In other words, the stage at which oxidative maturation became most evident in our system was not the one with the clearest signature of increased active PDH. Instead, the strongest late signal was the rise in PC together with LDHB, which points to increasing pyruvate carboxylation and therefore increasing anaplerotic support. That interpretation corroborates earlier work in iPSC-derived MNs (Vandoorne *et al*, 2019) and broader evidence that neural tissue depends on anaplerotic reactions to replenish TCA-cycle intermediates that are continuously withdrawn for biosynthesis and transmitter-related metabolism (Marin-Valencia *et al*, 2010; Hassel, 2001; Sonnewald & Rae, 2010). We cannot conclude from these data that PC simply replaced PDH in MNs, but the pattern of decreased PDHA1 combined with increased PC, LDHB, and pPDHA1 strongly argues that late MN maturation involves more than a straightforward increase in canonical pyruvate oxidation. Instead, mature MNs appear to reinforce oxidative metabolism with anaplerotic input, maybe to meet the unusual spatial and metabolic demands imposed by large somata, long axons, and specialized synaptic terminals. In that sense, the d14-to-d25 interval is best understood as the stage at which mitochondrial activation, mitochondrial remodeling, and carbon rerouting finally converge into a mature MN-like oxidative and largely anaplerotic phenotype.

Finally, the patterns we describe here may also matter beyond normal development. Mitochondrial dysfunction is a recurring theme in MN disease, including ALS and CMT-related disorders (van Lent *et al*, 2021; Mou *et al*, 2021; Vandoorne *et al*, 2018). In particular, Mfn2 dysfunction has been linked to impaired mitochondrial transport and abnormal MN biology (Mou *et al*, 2021; Filadi *et al*, 2015). The late stages of differentiation, when Mfn2- and Drp1-associated remodeling become prominent, may represent a point at which MNs become especially dependent on mitochondrial network organization. That could be relevant for interpreting disease phenotypes in iPSC-based MN models.

In conclusion, our data show that metabolic maturation during human iPSC-to-MN differentiation is directional, but not monotonic (Fig. 5B). Rather than occurring as a simple switch from glycolysis to oxidative phosphorylation, the transition unfolded in stages, with an early exit from the pluripotent metabolic state from stem cell to early neuroepithelial, a distinct mixed profile at a late neuroepithelial stage, a major step of mitochondrial activation at the transition from neuroepithelial to neural precursor, and a later maturation phase toward the maturing neuron in which oxidative metabolism became more extensively established (Fig. 5B). Importantly, this late phase was not defined by a simple increase in canonical PDH-linked pyruvate oxidation. Rather, the combined rise in PC, LDHB, and pPDHA1 pointed to a metabolic state in which oxidative metabolism relies less on PDH and more on anaplerotic input (Fig. 5B). Together, these findings suggest that MN maturation requires not only more active mitochondria, but also progressive remodeling of mitochondrial organization and more flexible routing of carbon into the TCA cycle. This stage-resolved view of metabolism throughout differentiation adds clarity to the current model of MN development and may also help to better understand why these cells are so vulnerable to mitochondrial dysfunction in disease.

## Methods

### Human Induced Pluripotent Stem Cell Culture

Cells were grown under feeder-free conditions using either mTESR1 or mTESRPlus cell culture medium in Geltrex-coated 6-well plates. For passaging, colonies were harvested using the enzyme-free passaging reagent, Versene, once they reached 60-80% confluency. The hiPSC line 028#1, kindly provided by Philipp Koch of the HITBR Hector Institute for Translational Brain Research, Mannheim, was used (Hörner *et al*, 2021; Couturier *et al*, 2024).

### Motoneuron Differentiation

MN differentiation was adapted from published protocols (Hörner *et al*, 2021; Du *et al*, 2015) with modifications to improve reproducibility. In brief, the process consisted of the following stages: (i) Neuroepithelial induction (d0–1): at 60–70 % confluency iPSCs were switched to MNDM1 medium (Neurobasal/DMEM-F12 with N2, B27, Glutamax, antibiotics, ascorbic acid, CHIR99021, DMH1, SB431542). Cells were replated on Geltrex with ROCK inhibitor for one day. (ii) Progenitor expansion (d2–12): Cells were maintained in MNDM1 until day 6, then passaged into MNDM2 medium (ascorbic acid, CHIR99021, DMH1, SB431542, purmorphamine, retinoic acid). (iii) Neurosphere formation (d12–18): MN progenitors were aggregated into neurospheres and cultured in MNDM3 medium (ascorbic acid, purmorphamine, retinoic acid, BDNF, GDNF, IGF-1). (iv) Terminal differentiation (d18 onward): Neurospheres were dissociated and plated on coated coverslips in MNDM4 medium (ascorbic acid, purmorphamine, retinoic acid, BDNF, GDNF, IGF-1, Compound E) for maturation. A full step-by-step protocol, including detailed medium compositions and catalog numbers, is provided in the Supplementary Materials (Text S2).

### Immunostaining and Fluorescence Imaging

Cells were washed once with PBS prior to incubating them with 4 % paraformaldehyde at room temperature for 15 minutes. Subsequently, cells were washed twice with an appropriate volume of PBS and stored in PBS at 4 °C until they were stained and imaged. For immunostaining, the cells were permeabilized with 0.1 % Triton X-100/PBS for 5 minutes at room temperature. They were then washed three times with Tris-buffered saline supplemented with 0.1 % TWEEN (TBST). Afterwards, 2 % BSA/PBS was added as a blocking solution and left for one hour at room temperature. During that time, the primary antibody solutions were prepared in 2 % BSA/PBS, and after one hour had passed, incubated with the cells at 4 °C overnight. The next day, cells were washed three times with TBST. Then the secondary antibody solution was added to the cells and left to incubate for 1 hour at room temperature. From here on out, the protocol was performed under dark conditions. When the incubation was over, cells were washed twice with TBST and twice with PBS. The cell culture vessel was then filled with PBS and placed in the microscopy room until the next day to equilibrate its temperature. Primary antibodies used and their dilution factors (DF) were as follows: Oct-4 (DF: 1:500), SOX2 (DF: 1:500), pPDHA1 (DF: 1:500), PDHA1 (DF: 1:200), LDH (DF: 1:600), LDHA (DF: 1:200), LDHB (DF: 1:200), Mfn1 (DF: 1:1000), Mfn2 (DF: 1:500), Drp1 (DF: 1:1000), PC (DF: 1:1000), TIGAR (DF: 1:500), Lamin A/C (DF: 1:1000), Hb-9 (DF: 1:100), Isl-1 (DF: 1:100), GAPDH (DF: 1:300), Tuj1 (DF: 1:500), vAChT (DF: 1:400). A complete list of all catalog numbers and suppliers may be found in the supplementary materials (Table S1). Fixed samples were imaged using a Leica DMi8 widefield fluorescence microscope equipped with a N PLAN L 20x/0.35 DRY objective and a Leica K5 sCMOS camera. Fluorophores were detected using the following filter cube configurations: DAPI (405/60 EX; 455 DC; 470/40 EM), GFP (470/40 EX; 495 DC; 525/50 EM), DSRED (546/10 EX; 560 DC; 605/75 EM), and Cy5 (635/18 EX; 652 DC; 680/42 EM). Images were acquired with exposure times between 3 and 200 ms, depending on channel.

### Mitotracker Mitochondrial Morphology Assay

At the start of the experiment, the cell culture medium was pre-warmed in a 37 °C water bath for 5 minutes. Nuclei were stained using Hoechst 33342. It was prepared at a stock concentration of 10 mg/mL and diluted in pre-warmed cell culture medium to a final concentration of 1 µg/mL. To stain the mitochondria, Mitotracker Red CMXRos was used. It was prepared at a stock solution of 1 mM in DMSO and diluted in pre-warmed cell culture medium to a final concentration of 200 nM. At the time of starting the experiment, cells were washed once with pre-warmed medium and then incubated with pre-warmed medium containing Hoechst and Mitotracker for 30 minutes at 37 °C with 5 % CO_2_-supplementation. Following incubation, three washes with pre-warmed medium were performed before beginning live-cell imaging. Live-cell imaging was performed using a Leica TCS SP8 confocal microscope with the following parameters: 37 °C with 90 % RH and 5 % CO_2_ supplementation, 512 x 512 pixels image dimension, 1400 Hz scan speed, 80 x 80 x 300 nm (x,y,z) voxel size with a 4.5 x zoom applied and pinhole set to 1 airy unit, 405 nm excitation laser with detector bands set to 410 - 539 nm, 552 nm excitation laser with detector bands set to 557 - 764 nm, bidirectional x scanning turned on, and using a HC PL APO CS2 63x/1.40 OIL objective.

### JC1 Mitochondrial Membrane Potential Assay

Cell culture growth medium, similar to the Mitotracker assay, was pre-warmed at 37 °C prior to beginning the experiment. Hoechst was also used at a final concentration of 1 µg/mL. 5,5′,6,6′-tetrachloro-1,1′,3,3′-tetraethylbenzimidazolocarbocyanine iodide (JC1) was prepared at a stock concentration of 1 mg/mL in DMSO. Final concentration in the cell culture growth medium was 0.5 µg/mL. Similar to the Mitotracker assay, cells were incubated with medium containing JC1 and Hoechst for 30 minutes at 37 °C with 5% CO_2_ supplementation. The same three washes were performed using pre-warmed medium prior to commencing the imaging session. Live-cell imaging was performed using a Leica TCS SP8 confocal microscope with the following parameters: 37 °C with 90 % RH and 5 % CO_2_ supplementation, 1024 x 1024 pixels image dimension, 600 Hz scan speed, 180 x 180 x 500 nm (x,y,z) voxel size and pinhole set to 1 airy unit, 405 nm excitation laser with detector bands set to 421 - 509 nm, 488 nm excitation laser for the simultaneous excitation of both free and aggregate forms of JC1 with detector bands set to 515 - 545 nm for the free-form of JC1 (green) and 572 - 608 nm for aggregate form of JC1 (red), bidirectional x scanning turned on, and using a HC PL APO CS2 63x/1.40 OIL objective.

### Extracellular Glucose Assay

Cells were plated in a Geltrex-coated 24-well plate. For specific cell numbers, please refer to the motoneuron differentiation protocol in the supplementary materials. Cell splits were performed on the stated days in the protocol. After 24 hours, the cell growth culture medium was changed to fresh medium, and cells were allowed to proliferate for another 24 hours. Subsequently, the supernatant was taken from 10 random wells to ensure representative sampling, centrifuged at 13,000 x g for 10 minutes to remove all cellular debris, and stored at -80 °C until samples were used for the experiment. For extracellular glucose measurement, a fluorimetric assay that measures D-glucose concentration was used. A modified version of the manufacturer’s protocol was used, where the assay buffer was replaced with an in-house buffer solution containing 25 mM HEPES and 150 mM NaCl, calibrated to a pH of 7.4. NaCl was supplemented to form an isotonic solution. Assays were performed in black fluorescence assay plates, and read-outs were measured using a CLARIOstar Plus plate reader (BMG Labtech). Relevant information about the kit, including catalog number and supplier can be found in the supplementary materials (Table S1).

### NMR-based extracellular metabolite analysis

Supernatant samples were diluted with D_2_O (Sigma Aldrich) and were transferred into 5 mm NMR tubes (Wilmad lab glas). For quantification, Trimethylsilylpropionic acid (TSP) was added to a final concentration of 0.5 mM in D_2_O. All 1H NMR measurements were carried out on a JEOL 14.1 T JMTC-600/54/XC/WS spectrometer (600 MHz for 1H). The 90° proton pulse was calibrated to 7 µs at 8.9 dB, corresponding to a nutation frequency of 35.7 kHz. Water was suppressed using a presaturation pulse sequence with the presaturation pulse power of 65 dB. 35,874 data points were acquired with 64 scans and 4 prescans with a relaxation delay of 4s. The NMR data were processed in Julia 1.12 with an NMR package developed by Marcel Utz (M.Utz, NMR Package for Julia Programming Language, 2021, https://github.com/marcel-utz/NMR.jl.git and cited in (Barker *et al*, 2026)). For spectral processing, the free induction decay (FID) signals were Fourier transformed and zero-filled to 2^17^ data points, and an apodization function with 0.5 Hz line broadening was applied. Zero-order phase correction and baseline correction were performed. Peak assignments were based on reference values from Chenomx and the Human Metabolome Database (HMDB) and are listed in Supplementary Table S2.

### Image Segmentation and Analysis

Mitotracker image analysis: Mitotracker images were deconvolved in post-processing using the Huygens deconvolution wizard (Scientific Volume Imaging) prior to analysis. For each condition, 50 randomly selected cells were manually outlined to generate cell masks. The mitotracker channel was subsequently used for 3D segmentation of individual mitochondria in Ilastik (Berg *et al*, 2019). Segmented masks were then loaded into Fiji and analyzed using the MorpholibJ plugin for 3D morphological feature extraction. Regarding immunostaining image analysis, for each condition, image sets from 3 independent experiments were loaded into Cellprofiler (Stirling *et al*, 2021). Background subtraction was made by the following method. First, the mean gray value of intensity (mean) and its standard deviation (std) were measured in an area containing cells of a negative control sample (only secondary antibodies, without primary antibody incubation). Then, the value “mean + 2*stdev” was subtracted from the entire image to be analysed. Following, a threshold was set to distinguish between the foreground and the background of the image, and out of this threshold, a mask was created. Finally, quantification was achieved by measuring the mean inside this mask in the original background-subtracted image. This process was completed for every channel of each raw multi-channel image. JC1 image analysis: For each condition, image sets from 3 independent experiments were loaded into Cellprofiler. A threshold was set in both the green and red channels to distinguish meaningful signal from the image background. The mean intensity and area occupied were thus measured in both channels. Finally, ratios of red:green intensity and area were calculated.

### Statistical Analysis and Graphing

All statistical analyses and data visualizations were performed in R (version 4.4.2; R Core Team, 2024) using the integrated development environment RStudio (version 2025.5.0.496; Posit team, 2025). Key packages included tidyverse (Wickham *et al*, 2019), dplyr (Wickham *et al*, 2014), and ggplot2 (Wickham, 2016). Normality was assessed where appropriate. Group comparisons were conducted using parametric or nonparametric tests. Parametric testing employed one-way ANOVA followed by Tukey’s post-hoc test, while nonparametric testing used Kruskal-Wallis with Wilcoxon post-hoc comparisons. If other statistical tests were used, it is further specified in the relevant figure legends. Statistical significance is indicated as follows: * *p* < 0.05; ** *p* < 0.01; *** *p* < 0.001; n.s = not significant.

## Data availability

Data will be made fully accessible upon request.

## Author contributions

Conceptualization, JJ and RR; Data curation, AR and JJ; Formal analysis, AR and JJ; Funding acquisition, MH and RR; Investigation, AR, JJ, RR; Methodology, AR, JJ; Project administration, RR; Resources, AR, MH, RR; Software, AR and JJ; Supervision, RR; Validation, AR and JJ; Visualization, AR and JJ; Writing – original draft, JJ and RR; Writing –review & editing, AR, JJ, MH, RR.

## Acknowledgements

JJ was supported by a fellowship of the graduate training group Perpharmance, (grant BW6_07, Baden-Württemberg Ministry of Science, Research and Arts), and is a current doctoral candidate within the Heidelberg Biosciences International Graduate School (HBIGS). RR was funded by DFG grant INST874/9-1.

## Supplementary

Fig. S1-S4, Tables S1 and S2, Texts S1 and S2 Mitochondrial rotation and masks video

